# A transparent multicriteria and fuzzy classification approach for genome-based probiotic candidate prioritisation

**DOI:** 10.64898/2026.07.20.739529

**Authors:** Nour ElHouda Ounissi, Mohamed Amine Gomri, Mohamed El Hadef El Okki

## Abstract

The identification of novel probiotic candidates with potential health-promoting properties remains a major challenge in food biotechnology and increasingly relies on in silico screening of genomic information. However, probiogenomic markers are heterogeneous, and safety, survival-colonisation, and functional-benefit traits do not contribute equally to probiotic potential. This study developed the Structured Probiotic Potential Index (SPPI), a fuzzy multicriteria system for genome-based probiotic candidate prioritisation. A hierarchical evaluation structure was established from probiogenomic evidence and organised into three main pillars and fourteen subcriteria. Expert judgements were collected using the Analytic Hierarchy Process, followed by consistency-based curation and weight aggregation. A curated dataset of 48 complete bacterial genomes, distributed into probiotic, potentially probiotic, neutral, and pathogenic groups, was taxonomically validated and analysed using an automated probiogenomic screening pipeline. Genome-wide screening generated 3,218 binary genomic features, from which curated probiogenomic markers were mapped to the scoring hierarchy. The resulting index was formulated as a normalised expert-weighted equation integrating safety, survival-colonisation, and functional-benefit components. SPPI prioritised genomes according to weighted probiogenomic profiles and separated pathogenic genomes from favourable probiotic and potentially probiotic profiles within the analysed dataset. Downstream Fuzzy Comprehensive Evaluation transformed the continuous score into probiotic/potentially probiotic, neutral, and pathogenic classes. Under internal leave-one-out evaluation, all 48 genomes were assigned to their expected reference classes, while confidence analysis distinguished high-confidence from borderline assignments. Post-classification comparison with ProbML showed concordant behaviour for most genomes and discordant predictions for selected cases. These results support SPPI as a transparent genome-based decision-support system for early probiotic candidate prioritisation before experimental validation.

## Introduction

Long before probiotics entered the genomic era, their scientific history began with empirical observations that specific microorganisms associated with fermented foods and the intestinal environment could influence host health. From the early work of Metchnikoff on fermented milk to Tissier’s observations on bifidobacteria and the subsequent development of commercial probiotic products, probiotics became central to food biotechnology, microbiome research, and health-related microbial applications (Siezen & Wilson, 2010; Valdez-Baez et al., 2022). Today, the field has expanded beyond traditional fermented matrices toward microbiome-based strategies, functional foods, and live biotherapeutic products, reflecting growing interest in microorganisms able to support host physiology through targeted biological functions (Castro-López et al., 2021; Churin et al., 2026; Pathan, Karlegan, & Shih, 2026).

Classical probiotic selection has relied heavily on phenotypic screening, including acid and bile tolerance, adhesion, antimicrobial activity, and safety evaluation. These assays remain essential, but they are time-consuming, strain-dependent, and difficult to scale across large microbial collections. The absence of universal phenotypic attributes shared by all probiotic strains has therefore encouraged the integration of genome-based approaches aimed at improving early candidate assessment (Castro-López et al., 2021; Choi et al., 2020). Probiogenomics offers a rapid and reproducible means to explore safety determinants, survival-colonisation traits, host-interaction markers, metabolic capacities, and bioactive potential directly from whole-genome data (Carpi et al., 2022; Khullar et al., 2022; Najar et al., 2025; Sabino et al., 2025).

Despite these advances, interpreting probiogenomic data remains challenging. Many genomic markers considered favourable for probiotic assessment are not specific to probiotic strains and may also occur in commensal, environmental, opportunistic, or pathogenic bacteria. Conversely, safety-associated determinants require contextual interpretation because their biological meaning depends on genomic localisation, mobility, sequence identity, completeness, and expression potential. Recent genomic resources and machine-learning approaches have improved the exploration of probiotic potential, yet their outputs may remain difficult to interpret when heterogeneous biological signals are reduced to a single label (Neres Rodrigues et al., 2026; Orkkatteri Krishnan et al., 2025).

A major methodological need is therefore the development of transparent systems able to integrate heterogeneous genomic evidence according to its biological relevance. The Analytic Hierarchy Process (AHP) provides a structured expert-based method for deriving relative weights from pairwise comparisons, whereas Fuzzy Comprehensive Evaluation (FCE) enables multicriteria class assignment when biological boundaries are gradual and not strictly binary (Saaty, 1987; Saaty, 1977, 1984; Yang et al., 2025; Zadeh, 1965). Such a system should not treat all detected markers as equivalent, but should organise them according to their biological role in probiotic candidate evaluation. Combining these approaches with automated probiogenomic screening can support a prioritisation logic in which biosafety is evaluated first, while favourable survival-colonisation and functional traits refine the ranking of candidates with acceptable safety profiles.

This study aimed to develop SPPI (Structured Probiotic Potential Index), an AHP-guided fuzzy multicriteria system for the in silico prediction and prioritisation of probiotic potential across bacterial genomes. The workflow combines literature-guided marker selection, expert-derived weighting, automated genome screening, normalised scoring, FCE-based class assignment, and post-classification comparison with ProbML as an external reference tool. The objective was not to replace experimental validation, but to provide an interpretable genome-based decision-support layer for early candidate selection.

## Materials and methods

### Study design and methodological workflow

This study followed a stepwise computational design to construct and evaluate SPPI (Structured Probiotic Potential Index) as a genome-based multicriteria scoring system. The workflow integrated literature-guided probiogenomic criteria selection, hierarchical multicriteria structuring, expert-based weighting using the Analytic Hierarchy Process (AHP), curated genome dataset construction, taxonomic confirmation, automated probiogenomic screening, weighted score calculation, and downstream Fuzzy Comprehensive Evaluation (FCE) classification. The complete methodological workflow is summarised in Fig. 1.

**Fig. 1.**
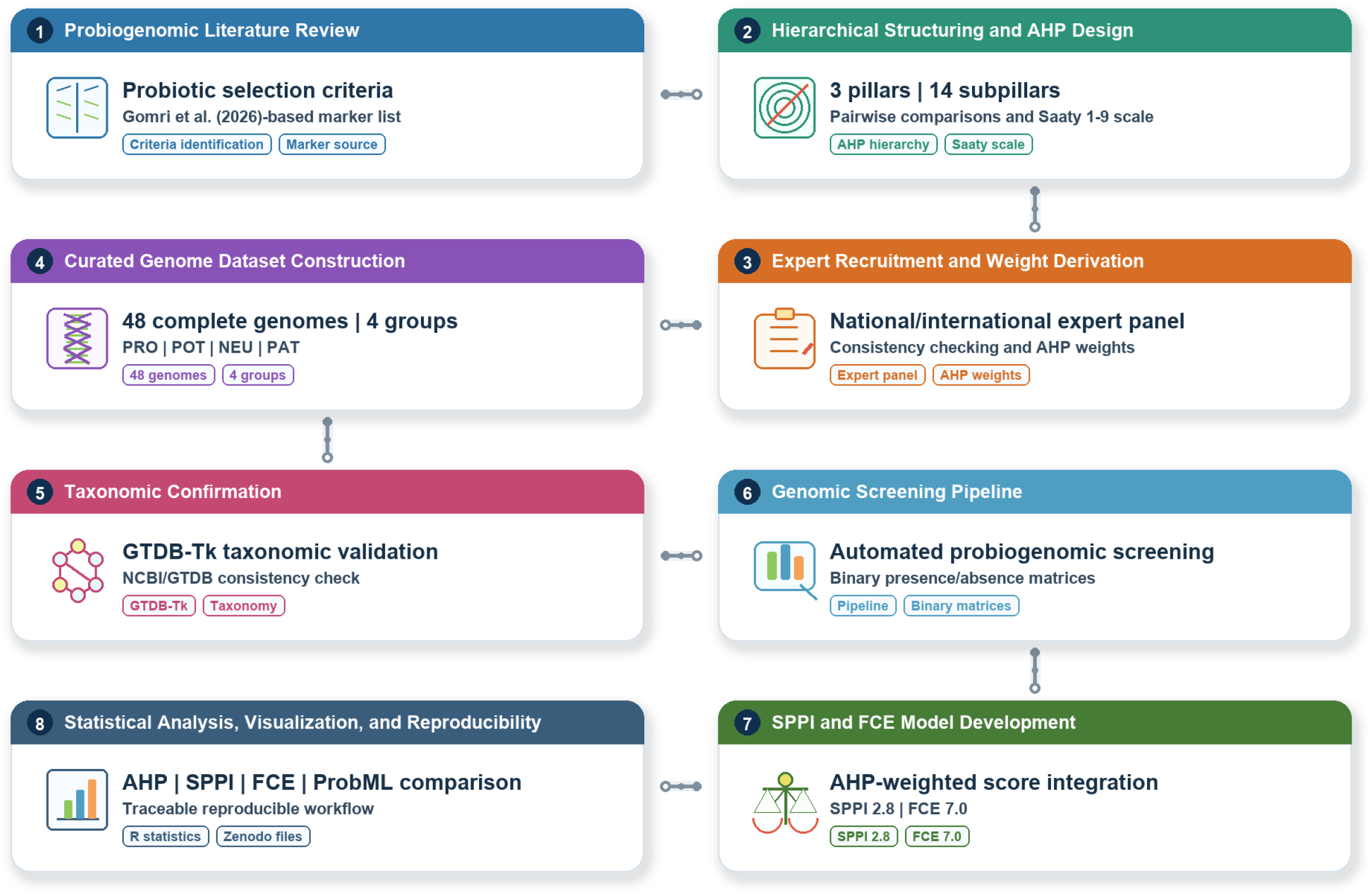
Methodological workflow for the development of an in silico probiotic scoring system from prokaryotic genomes.

### Literature-guided criteria selection and hierarchical structuring

A targeted review of the probiogenomic and comparative genomics literature was conducted to identify genomic traits relevant to the in silico evaluation of probiotic potential. The retained criteria covered three complementary dimensions of candidate assessment: biosafety, survival-colonisation, and functional and metabolic benefits. These dimensions were selected because probiotic potential cannot be inferred from a single genomic determinant, but rather depends on the combined interpretation of safety-related constraints, adaptation to gastrointestinal conditions, host-interaction traits, and potential functional activities (Castro-López et al., 2021; Khullar et al., 2022; Sabino et al., 2025).

The genomic marker framework was primarily derived from Gomri et al. (2026). Related probiogenomic and comparative genomic studies supported the identification of the broader probiotic selection criteria used to construct the hierarchical structure (Carpi et al., 2022; Najar et al., 2025; Sabino et al., 2025). However, the retained markers were not interpreted as probiotic-specific determinants, as several correspond to conserved bacterial functions that may also occur in commensal, environmental, or pathogenic bacteria. Consequently, detected markers were interpreted as components of a composite genomic profile during downstream scoring (Neres Rodrigues et al., 2026; Papadimitriou et al., 2015). An overview of the hierarchical criteria, marker groups, and scoring contribution used for SPPI construction is provided in Table 1.

**Table 1.**
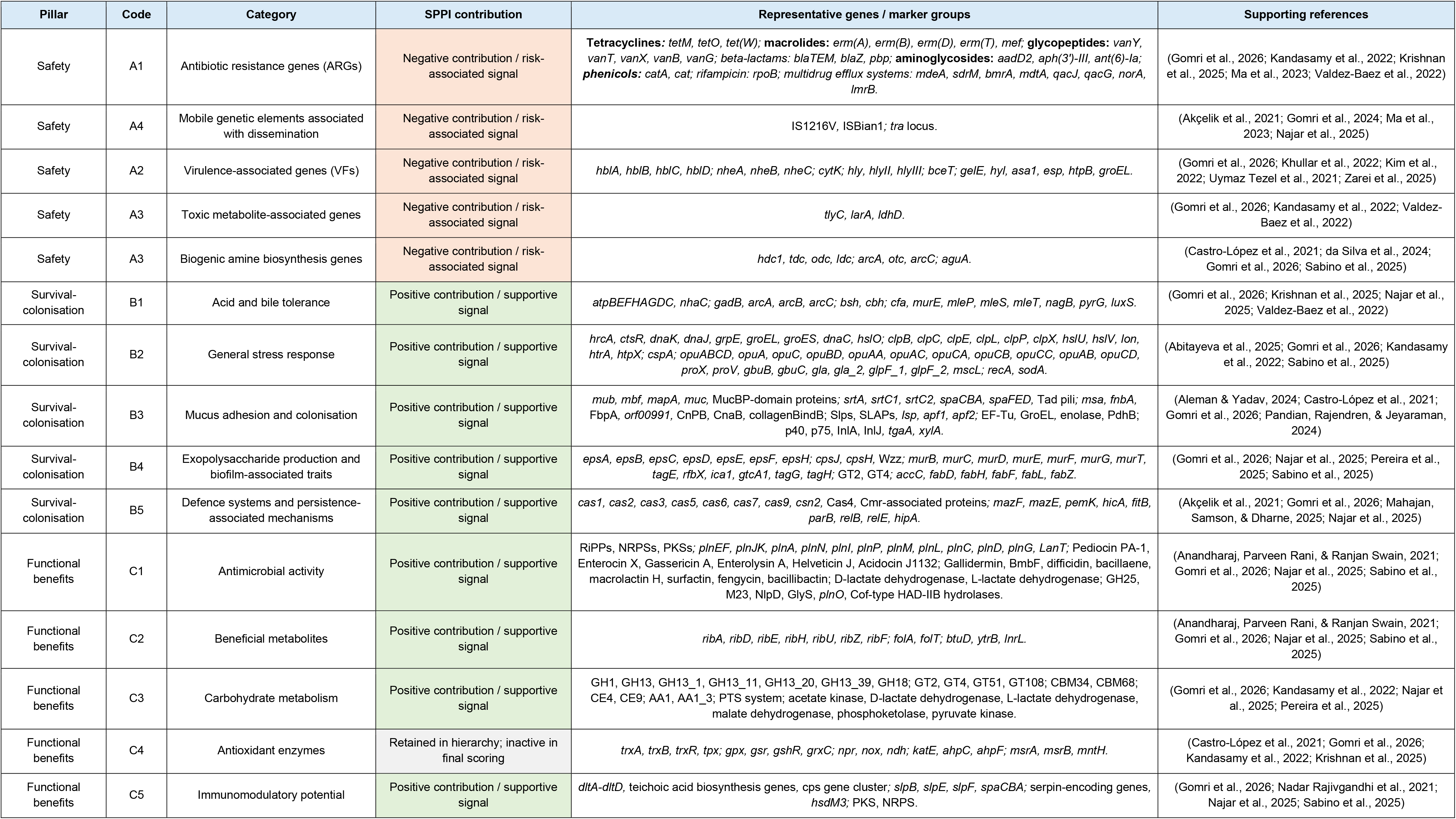
Integrated hierarchical marker framework used for SPPI construction.

The retained criteria were thus organised into three main pillars and fourteen subpillars. Biosafety included antimicrobial resistance, virulence-associated determinants, toxin-related signals, and genomic instability. Survival-colonisation included acid and bile tolerance, general stress response, mucus adhesion and persistence, exopolysaccharide production, and defence systems. Functional and metabolic benefits included antimicrobial activity, beneficial metabolites, carbohydrate metabolism, antioxidant enzymes, and immunomodulatory potential.

### AHP questionnaire design, expert recruitment and weight derivation

AHP was used to derive relative importance weights for the probiogenomic evaluation criteria through structured pairwise comparisons (Saaty, 1987; Saaty, 1977, 1984). The hierarchy generated one 3 × 3 matrix for the three main pillars, one 4 × 4 matrix for the Safety subpillars, and two 5 × 5 matrices for the Survival-colonisation and Functional and Metabolic benefits subpillars. The AHP hierarchical levels, decision components, included criteria, and corresponding pairwise comparison matrix sizes are summarised in Table 2. Pairwise comparisons were performed using the Saaty 1-9 scale, where 1 indicates equal importance and higher values indicate increasing preference of one criterion over another; reciprocal values represented inverse judgements (Fig. 2).

**Fig. 2.**
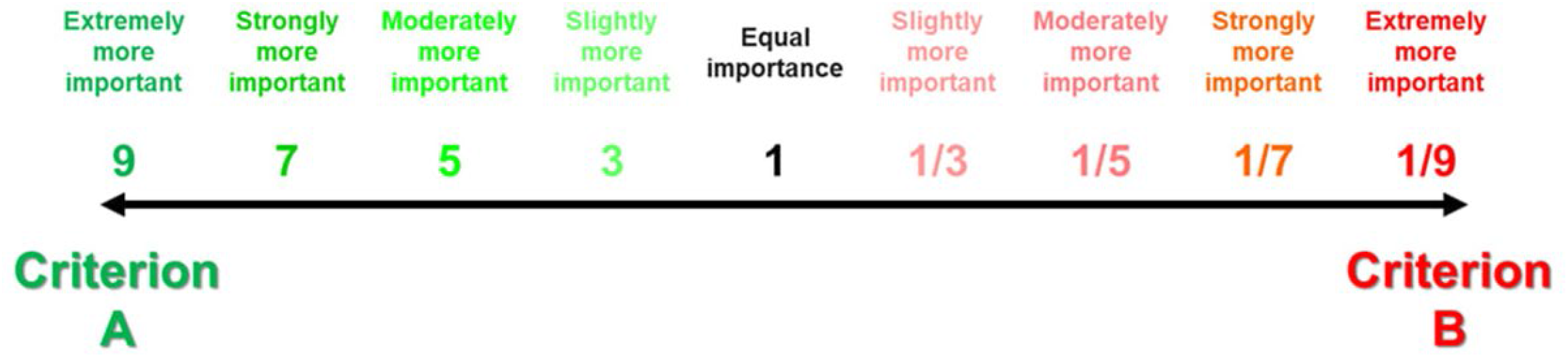
Saaty scale used for the pairwise comparison of probiotic criteria. The expression 1/n indicates that a value of n is assigned to criterion B when criterion A is assigned a value of 1.

**Table 2.** AHP hierarchy and pairwise comparison matrices used in the study.

| Hierarchical level | Decision component | Elements included | Matrix size |
| --- | --- | --- | --- |
| **Level 0** | Goal | Evaluation of probiotic genomic potential | Not Applicable |
| **Level 1** | Main pillars | Safety; Survival and Colonisation; Functional and Metabolic Benefits | $3 \times 3$ |
| **Level 2A** | Safety subpillars | A1 Antimicrobial resistance; A2 Virulence; A3 Toxin production; A4 Genomic instability | $4 \times 4$ |
| **Level 2B** | Survival and Colonisation subpillars | B1 Acid and bile tolerance; B2 General stress response; B3 Mucus adhesion capacity; B4 Exopolysaccharide production; B5 Defence systems | $5 \times 5$ |
| **Level 2C** | Functional and Metabolic Benefits subpillars | C1 Antimicrobial activity; C2 Beneficial metabolites; C3 Carbohydrate metabolism; C4 Antioxidant enzymes; C5 Immunomodulatory potential | $5 \times 5$ |

The questionnaire was implemented using Google Forms and included 38 structured questions corresponding to the different levels of the hierarchy. Before dissemination, it was pre-tested by a multidisciplinary internal expert panel to verify question clarity, logical organisation, scale interpretation, and consistency between questionnaire items and the hierarchical structure. Experts were then recruited through bibliographic screening in probiotics, microbiology, biotechnology, genomics, bioinformatics, and food safety. In total, 516 national and international experts were included in the contacted/referred pool, and 41 complete responses were retained at questionnaire closure (Supplementary Table S1).

Google Forms responses were exported to spreadsheet format and processed using an automated AHP calculator. The calculator reconstructed reciprocal pairwise comparison matrices for each expert and each hierarchical block, then computed priority vectors using the principal eigenvector method. For a set of n criteria, the reciprocal comparison matrix A = (aij) was defined such that aij expresses the relative importance of criterion i over criterion j and aji = 1/aij. The priority vector w was obtained from:

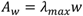

Where *λ_m_*_a*x*_ is the principal eigenvalue of the comparison matrix. Matrix consistency was evaluated using the consistency index and consistency ratio:

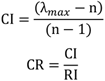

where RI is the random index corresponding to matrix size (Alonso & Lamata, 2006). CR values were interpreted at the matrix level. Matrices with CR ≤ 0.10 were considered acceptable, matrices with 0.10 < CR ≤ 0.20 were retained as moderately inconsistent but still usable for aggregation, and matrices with CR > 0.20 were excluded from the curated aggregation step (Apostolou & Hassell, 1993; Hussain Sinjar, 2017; Saaty, 2008). The 0.20 threshold was adopted because the questionnaire contained several independent comparison blocks and because moderate inconsistency may occur in complex expert-based biological judgements without invalidating all information provided by a respondent.

Curated aggregation was performed separately for each hierarchical section using only the matrices that satisfied the CR ≤ 0.20 criterion in that section. Aggregated matrices were generated by aggregating individual judgements using the geometric mean, followed by exact eigenvector-based weight calculation. This section-level curation preserved coherent expert information while limiting the influence of highly inconsistent pairwise comparisons.

### Curated genome dataset construction and taxonomic confirmation

A curated high-quality dataset of 48 complete bacterial genomes was constructed as a controlled reference panel for probiogenomic screening, SPPI development, and downstream model evaluation. Low-quality assemblies, duplicated entries, or poorly documented strains may bias functional interpretation; therefore, genome availability, strain-level traceability, and quality-based filtering were considered before downstream analysis (Gomri et al., 2026). Genomes were retrieved from the NCBI RefSeq Assembly database (https://www.ncbi.nlm.nih.gov/datasets/genome/) in March 2026 (Goldfarb et al., 2025; Sayers et al., 2026) and organised into four groups of twelve genomes each.

The first group, coded PRO, included clinically confirmed and/or commercially used probiotic strains, mainly human-associated strains. These strains were identified using Probio-Ichnos (https://probio-ichnos.streamlit.app/), MicrobiomePost (https://microbiomepost.com/), and supporting scientific literature (Petraro et al., 2024; Tsifintaris et al., 2024). The second group, coded PAT, included strains with documented pathogenicity in humans. These strains were selected from published scientific literature and reference pathogenic genome records available in NCBI RefSeq. The third group, coded NEU, included environmental or commensal strains used as an intermediate calibration group between favourable probiotic-related profiles and pathogenic profiles. These strains were selected from scientific literature and NCBI RefSeq records to include taxonomically diverse representatives with no documented probiotic use or human pathogenicity, allowing the scoring system to distinguish broadly distributed adaptive or metabolic traits from trait combinations more specifically informative for probiotic prioritisation. The fourth group, coded POT, included strains described as having probiotic potential based on in vitro evidence, without clinical validation or commercial use. These strains were identified using Probio-Ichnos and published probiotic screening studies, provided that complete RefSeq genome assemblies were available.

Only genomes with complete RefSeq assemblies, valid GCF accession numbers, genome completeness ≥ 90%, and contamination ≤ 5% were retained. Completeness and contamination values were evaluated using CheckM values reported in NCBI RefSeq entries (Parks et al., 2015). Genomes meeting all criteria were progressively selected until twelve genomes were retained for each group, while maintaining taxonomic diversity across genera and phyla.

For reproducibility and traceability, all retained genomes were organised using standardised folder and file names. Each genome folder included the group code, numerical identifier, genus, species, and strain information, whereas sequence and annotation files were renamed using short standardised genome codes, such as NEU01.fna, NEU01.faa, NEU01.gff, and NEU01.gbff. A metadata spreadsheet was also created to record genome codes, file names, BioProject identifiers, BioSample identifiers, RefSeq accessions, assembly metrics, isolation source, evidence type, and supporting references. The complete genome list and associated metadata are provided as Supplementary Table S2.

Taxonomic confirmation was performed using GTDB-Tk v2.3.2 against GTDB release R214 within the KBase environment (https://www.kbase.us/; accessed March 2026) (Arkin et al., 2018; Chaumeil et al., 2020, 2022).

GTDB-Tk classifications were compared with NCBI annotations to verify genus- and species-level consistency.

This validation step confirmed the taxonomic affiliation of the 48 retained genomes, and the validated assignments were retained as reference identifiers for downstream genomic screening and SPPI implementation.

### Automated probiogenomic screening pipeline

Following taxonomic confirmation, all retained genomes were analysed using a local modular probiogenomic screening pipeline (version 3.0) developed in a Linux Mint 22.2 environment. The pipeline transformed genome assemblies into structured binary outputs describing the detection or non-detection of genome-wide functional features and curated probiogenomic markers. Genome assemblies in FASTA format (.fna) were used as the only input files, and each genome was processed independently to maintain traceability of results. The complete workflow was executed in full analysis mode and required approximately 18 h for the 48-genome dataset, corresponding to about 22.5 min per genome, under the computational conditions used in this study. The pipeline was executed locally on a Dell Inspiron 15-3567 running Linux Mint 22.2, equipped with an Intel Core i5-7200U processor and 8 GiB of RAM. The modular workflow is summarised in Fig. 3.

**Fig. 3.**
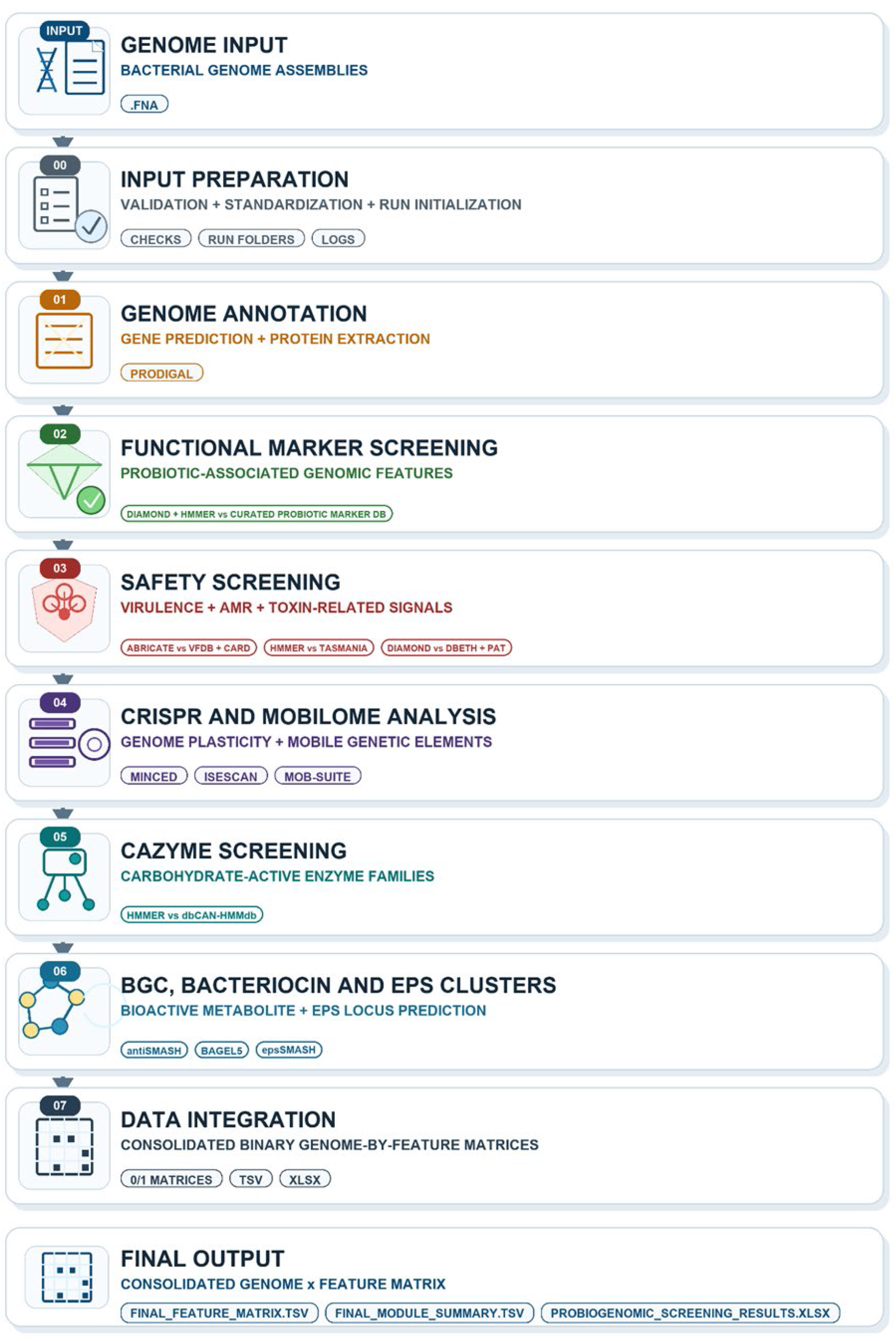
Overview of the modular bioinformatics pipeline used for probiogenomic screening of bacterial genomes.

For each genome, screened features were encoded as binary variables, with 1 indicating detection and 0 indicating non-detection under the implemented parameters. A value of 0 was therefore interpreted as absence of detectable evidence under the applied databases, thresholds, and tool versions, and not as definitive biological absence. This binary representation enabled construction of genome-by-feature matrices and marker-level files compatible with downstream mapping to the SPPI hierarchy.

The workflow integrated gene prediction and protein extraction, functional marker screening, virulence and antimicrobial resistance screening, toxin-related signal detection, CRISPR and mobile genetic element analysis, carbohydrate-active enzyme detection, and bioactive cluster screening. It included Prodigal, DIAMOND, HMMER, ABRicate-based screening against VFDB and CARD, TASmania, DBETH, PAT, MOB-suite, MINCED, ISEScan, dbCAN-HMMdb, antiSMASH, BAGEL5, and epsSMASH. Selected outputs, particularly those derived from antiSMASH, VFDB, and CAZyme analyses, were curated to retain features that could be directly interpreted within the defined scoring criteria and assigned to the corresponding SPPI pillars and subpillars. Final outputs consisted of consolidated binary matrices, marker-level presence/absence files, module summaries, and screening reports.

### SPPI 2.8 calculation

The binary outputs generated by the probiogenomic screening pipeline were mapped to the hierarchical SPPI structure. The pipeline produced a genome-by-feature matrix comprising 3,218 detected genomic features across the 48 genomes, encoded as binary variables. A feature-level mapping table was then used to assign 1,720 mapped feature entries to their corresponding SPPI subpillars. Of these entries, 1,447 were retained as active scoring features, whereas 273 were excluded from the scoring calculation. Across the mapping table, entries were classified as core, auxiliary, or excluded according to their role in the calculation. The scoring system was implemented using AHP-derived weights obtained from the expert-based prioritisation step.

The framework was developed through successive controlled iterations. Version 2.3 represented the first stabilised implementation of the hierarchical organisation, feature mapping, and calculation rules. Subsequent refinements were introduced in later iterations while preserving the original three-pillar and fourteen-subpillar structure. The retained 2.8 configuration was used as the final upstream scoring model.

For each genome *g*, safety-related evidence was first calculated as a risk-associated contribution:

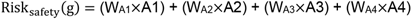

where A1, A2, A3, and A4 correspond to antimicrobial resistance, virulence-associated determinants, toxin-related signals, and genomic instability, respectively. The corresponding safety score was obtained by reversing the risk contribution:

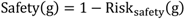

Survival-colonisation and Functional and Metabolic benefits were calculated as positive weighted components:

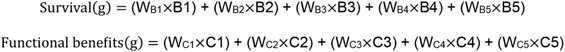

The final SPPI value was calculated as:

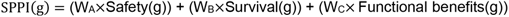

In this formulation, WA, WB, and WC represent the AHP-derived weights of the three main pillars. The final score was normalised from 0 to 1 and interpreted as a comparative genomic indicator of probiotic potential, not as direct experimental evidence of probiotic efficacy. Values closer to 1 correspond to profiles with stronger favourable genomic evidence and lower safety-associated constraints, whereas values closer to 0 correspond to profiles dominated by safety-related signals. The mapping files, scoring scripts, and generated score outputs were retained for reproducibility.

### FCE 7.0 classification and post-classification ProbML comparison

Following SPPI calculation, FCE was used to transform continuous scoring outputs into operational genome classes (Ma et al., 2023; Yang et al., 2025; Zadeh, 1965). Consequently, the four initial genome groups were reorganised into three categories: Category 1 for probiotic and potentially probiotic genomes, Category 2 for neutral genomes, and Category 3 for pathogenic genomes.

FCE Iteration 6.0 served as the reference downstream configuration. This configuration followed a two-stage hierarchical logic, in which safety-related descriptors were first used to identify Category 3 genomes, followed by a second stage designed to separate Category 1 from Category 2 among the remaining genomes. Based on this reference logic, FCE Iteration 7.0 was subsequently retained as the final downstream configuration.

This retained downstream configuration used the standardised outputs of SPPI 2.8 without modifying the upstream scoring algorithm. The first stage evaluated assignment to Category 3 using safety-related descriptors, including A2, A1, A3, and the aggregate Safety value. Genomes assigned to Category 3 were not processed further. The second stage was then applied to the remaining genomes and used SPPI together with Functional and Metabolic benefits, C2, and B3 to differentiate Category 1 from Category 2. Final class assignment was based on the highest membership degree.

ProbML was used only after FCE classification as an external reference tool for post-classification comparison. Genome sequences from the same 48-genome dataset were provided as FASTA input, and ProbML was executed locally through the MLG Dashboard environment (Orkkatteri Krishnan et al., 2025). Recorded outputs included prediction label, score, score distribution, model used, analysis date, and reference label. ProbML predictions were not used to define FCE membership functions, optimize FCE parameters, or assign FCE classes; they were used only to compare binary prediction behaviour on the same dataset after completion of the SPPI-FCE workflow.

### Statistical analysis, visualisation and reproducibility

Statistical analyses were conducted to support interpretation of the AHP weighting outputs, probiogenomic screening matrices, SPPI 2.8 values, and FCE 7.0 classification results. Analyses were performed using R v4.5.3 and RStudio v2026.01.2+418, while Python v3.11.9 was used for data structuring, score calculation, and workflow execution. Main R packages included ggplot2, vegan, pheatmap, and additional packages for statistical testing and visualisation (Kolde, 2025; Oksanen et al., 2026; Posit team, 2025; Wickham, 2016).

Expert-response data were summarised descriptively, and AHP consistency was evaluated through CR values calculated for each comparison matrix. For screening outputs, binary presence/absence matrices were used to quantify feature and marker richness across genome groups. Group-level differences were evaluated using Kruskal-Wallis tests followed, where appropriate, by Wilcoxon-Mann-Whitney pairwise comparisons with Benjamini-Hochberg correction (Benjamini & Hochberg, 1995; Kruskal & Wallis, 1952; Mann & Whitney, 1947). Fisher’s exact test was used to identify markers associated with specific genome groups (Fisher, 1922).

Because screening matrices were binary, Jaccard dissimilarity was used for genome comparisons. Principal coordinates analysis (PCoA), non-metric multidimensional scaling (NMDS), PERMANOVA, and beta-dispersion analysis were applied to examine multivariate organisation and group separation, using 9,999 permutations for permutation-based tests (Anderson, 2001, 2006; Gower, 1966; Jaccard, 1901; Kruskal, 1964). SPPI 2.8 outputs were compared across groups using Kruskal-Wallis tests, Welch’s ANOVA, Benjamini-Hochberg-adjusted pairwise comparisons, and Cliff’s delta effect sizes (Cliff, 1993; Welch, 1951). Principal Component Analysis (PCA) was applied to subpillar score profiles (Jolliffe & Cadima, 2016).

The retained FCE 7.0 model was evaluated using leave-one-out cross-validation, confusion matrices, balanced accuracy, precision, recall, and F1-score (Brodersen et al., 2010; Sokolova & Lapalme, 2009; Stone, 1974). For post-classification comparison with ProbML, FCE outputs were transformed into a binary structure corresponding to Category 1 versus non-Category 1. Binary performance was assessed using accuracy, sensitivity, specificity, Wilson confidence intervals, McNemar’s test, and Cohen’s kappa (Cohen, 1960; McNemar, 1947; Wilson, 1927). Reproducibility and traceability were supported by preserving genome metadata, binary matrices, AHP-derived weights, SPPI 2.8 files, FCE outputs, ProbML outputs, figures, and all associated Python and R scripts with the study data and code.

## Results

### Expert panel curation and AHP-derived weight structure

The AHP survey generated 41 complete expert responses retained at questionnaire closure from a contacted/referred pool of 516 experts. The contacted pool included international, national, and Scopus-identified experts, providing a geographically and academically diverse basis for expert-derived weighting. Because the questionnaire included several independent comparison blocks, consistency was evaluated separately for each AHP matrix. Using the retained CR ≤ 0.20 threshold, 18 of 41 pillar-comparison matrices were kept for aggregation. The Safety, Survival-colonisation, and Functional benefits blocks retained 31, 26, and 30 matrices, respectively. This matrix-level curation allowed the final weighting structure to be derived from the coherent part of the expert dataset while reducing the influence of highly inconsistent pairwise judgements.

After consistency-based curation and geometric-mean aggregation, the retained collective matrices showed improved coherence. At the pillar level, the aggregated consistency ratio decreased from 0.076 before curation to 0.013 after curation. The final pillar weights assigned the dominant contribution to Safety (73.6%), followed by Survival-colonisation (15.1%) and Functional benefits (11.4%) (Fig. 4A).

**Fig. 4.**
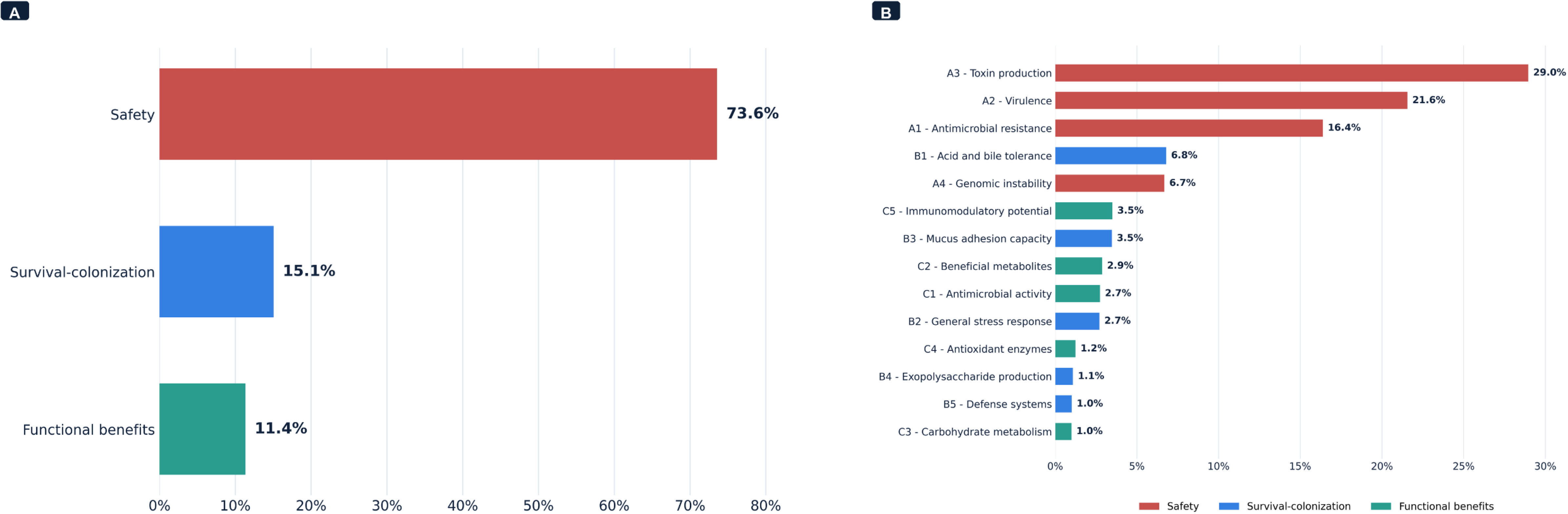
AHP-derived relative contributions to the final SPPI structure. A, relative contribution of the main pillars after consistency-based curation. B, relative contribution of the genomic subcriteria to the final SPPI structure.

At the subpillar level, toxin-related signals (A3, 29.0%), virulence-associated determinants (A2, 21.6%), and antimicrobial resistance contribution (A1, 16.4%) represented the three strongest global contributors (Fig. 4B). Together, these three safety-related components accounted for 67.0% of the final SPPI structure. The highest positive contribution outside Safety was assigned to acid and bile tolerance (B1, 6.8%), followed by immunomodulatory potential (C5, 3.5%), mucus adhesion and persistence (B3, 3.5%), and beneficial metabolites (C2, 2.9%).

The retained AHP weights were then integrated into the final SPPI 2.8 equations:

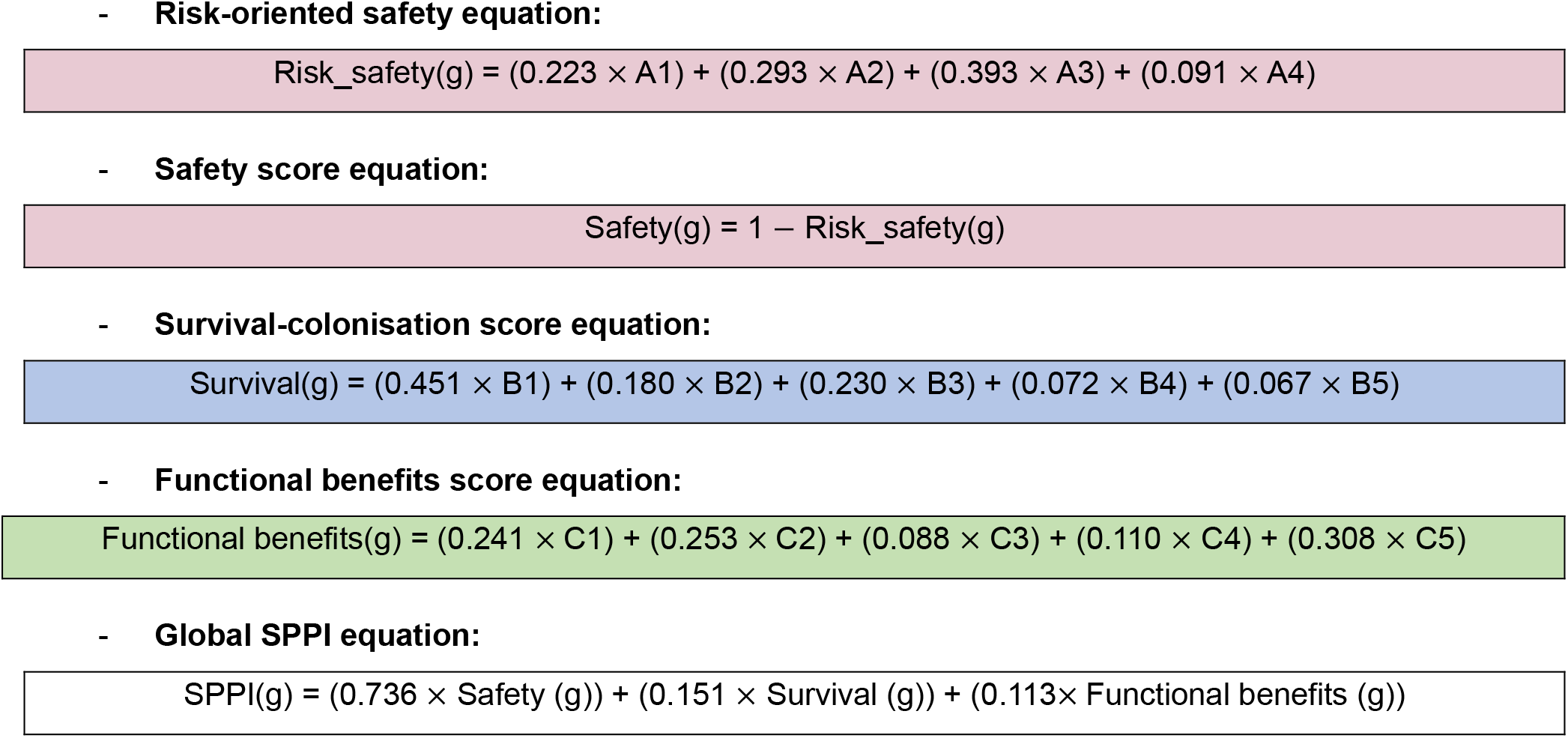

Here, *g* denotes the genome under evaluation, and A1-A4, B1-B5, and C1-C5 refer to the subcriteria defined in the scoring structure. These equations show that SPPI 2.8 was not calculated as a simple count of detected markers, but as a weighted genomic decision score integrating risk-associated and favourable probiogenomic evidence.

### Phylotaxogenomic validation of the curated genome dataset

Phylotaxogenomic validation using GTDB-Tk v2.3.2 against the Genome Taxonomy Database (GTDB release R214) within the KBase platform confirmed the taxonomic coherence of the curated genome dataset (Arkin et al., 2018; Chaumeil et al., 2020, 2022) (Fig. 5). All retained genomes were assigned to the expected taxonomic groups, with no conflicting affiliation between the initial NCBI RefSeq annotations and the GTDB-based classification. Consequently, no genome was reclassified or excluded at this stage.

**Fig. 5.**
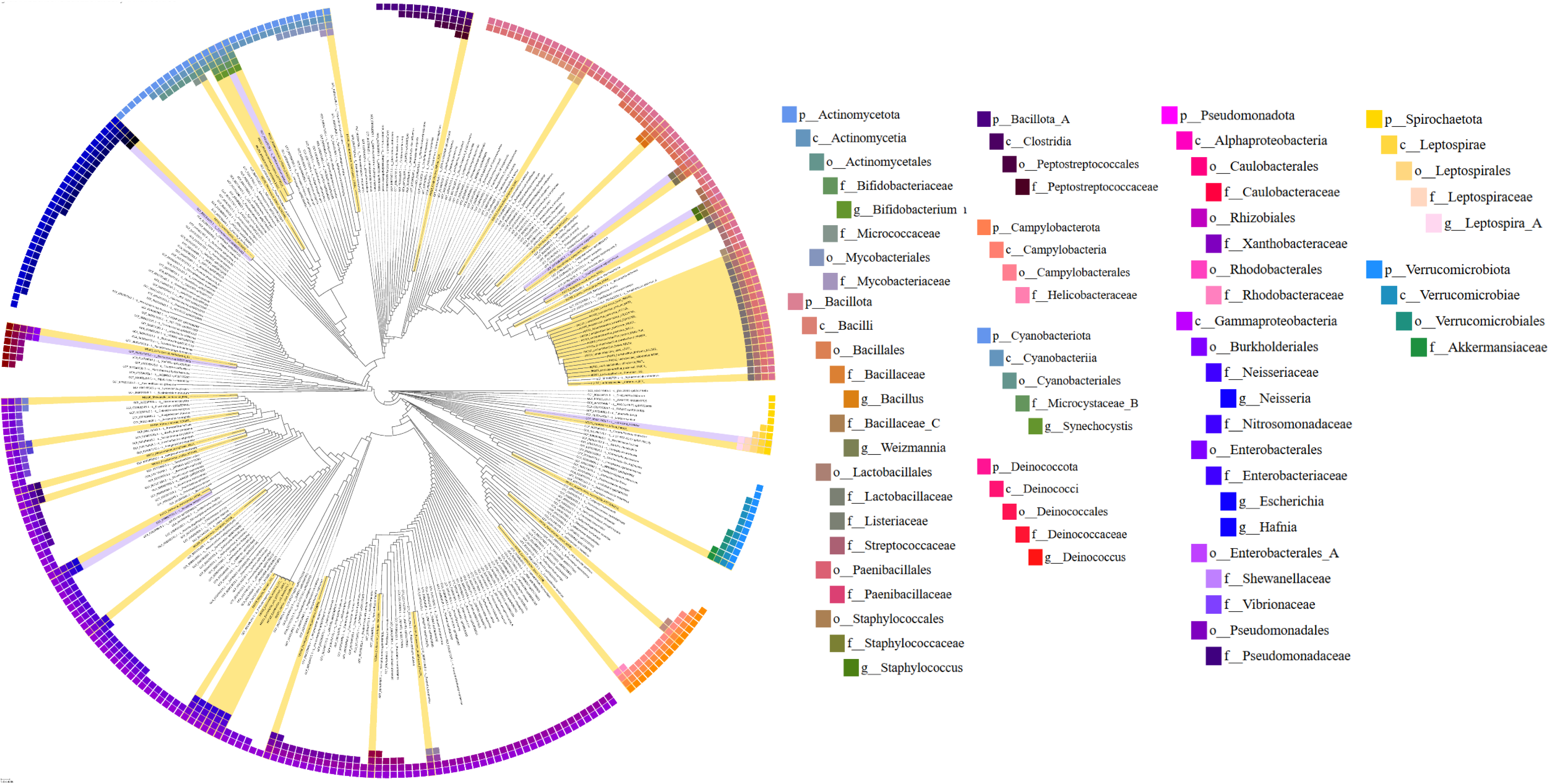
Phylotaxogenomic validation of the analysed genome dataset using Genome Taxonomy Database (GTDB) classification. Circular phylogenomic tree generated with Classify Microbes using GTDB-Tk v2.3.2, showing the taxonomic placement of all genomes; genomes analysed in this study are highlighted in yellow, confirming their expected affiliations and taxonomic coherence.

The phylogenomic tree showed that the curated dataset included genomes from several bacterial phyla, including *Actinomycetota*, *Bacillota*, *Campylobacterota*, *Cyanobacteriota*, *Deinococcota*, *Pseudomonadota*, *Spirochaetota*, and *Verrucomicrobiota* (Fig. 5). This taxonomic breadth provided a broader reference background for probiogenomic screening, scoring, and downstream classification, and reduced the likelihood that marker detection or model calibration would be driven mainly by lineage-specific genomic patterns.

This point is relevant because genomic signatures detected in heterogeneous bacterial datasets may reflect taxonomic background, ecological adaptation, or genome architecture, and do not constitute direct evidence of probiotic functionality (Neres Rodrigues et al., 2026). This interpretation is also consistent with Gomri et al. (2026), who reported a gap between probiotic genomic potential and experimental attention and suggested that current validation practices may contribute to survivorship bias by focusing on a limited set of canonical traits.

### Probiogenomic screening pipeline outputs

Following phylotaxogenomic validation, the probiogenomic screening pipeline generated a genome-wide functional feature matrix comprising 3,218 binary genomic features across the 48 genomes. These outputs were generated from the screening of 14 functional modules, including safety-related signals, accessory-genome features, mobilome-associated elements, carbohydrate-active enzyme families, bioactive clusters, exopolysaccharide-associated features, and curated markers selected for probiotic potential assessment. This matrix represented the complete automated screening output obtained before curated marker extraction and SPPI implementation.

At the genome-wide level, total feature richness differed significantly among the four genome groups (Kruskal-Wallis test, H = 8.46, df = 3, p = 0.037) (Fig. 6A). Pathogenic genomes displayed the highest feature counts, followed by neutral, probiotic, and potentially probiotic genomes. However, post hoc pairwise comparisons did not remain significant after multiple-testing correction, indicating that global feature richness alone was not sufficient to discriminate all groups. This pattern is consistent with the structure of the uncurated matrix, which included several safety-centred and accessory-genome modules. In particular, pathogenic bacteria may differ in virulence-factor content because accessory genomic elements can reshape genome architecture and modify virulence repertoires (Jackson et al., 2011), while antimicrobial-resistance determinants and mobile genetic elements may also contribute to group-level genomic differences (Cross, Partridge, & Sheppard, 2026; Liu et al., 2025).

**Fig. 6.**
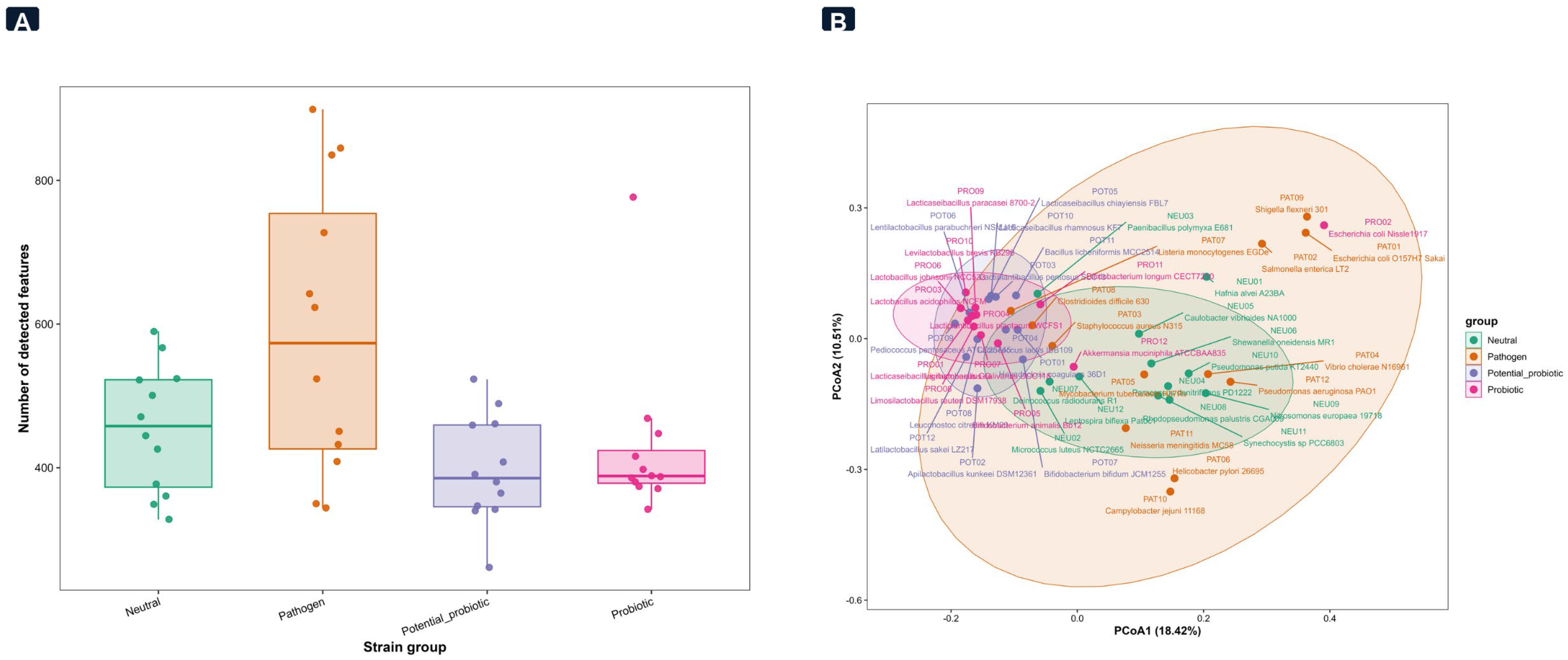
Genome-wide functional feature richness and multivariate structure across the four genome groups. A, total number of detected genomic features after genome-wide screening. B, multivariate structure of genome-wide binary feature profiles based on Jaccard distance and principal coordinates analysis (PCoA).

The genome-wide binary matrix was then examined using Jaccard-based PCoA (Fig. 6B). The first two coordinates explained 31.08% of the total variation, with PCoA1 accounting for 20.87% and PCoA2 for 10.21%. PERMANOVA detected a significant group effect on feature composition (R² = 0.165, F = 2.90, p < 0.001). The moderate R² value indicates that genome group explained part of the observed variation, while a substantial fraction remained genome-specific. Thus, genome-wide feature composition showed partial group separation, but also reflected the broader heterogeneity expected in comparative bacterial genomics (Bhattacharya, Joishy, & Khan, 2025; Can et al., 2025).

### Curated marker composition and marker-level separation

The curated marker analysis included 203 markers from the probiogenomic marker framework reported by Gomri et al. (2026), grouped into 13 main functional categories and evaluated across the same 48 genomes. The total number of curated markers did not differ significantly among the four genome groups (Kruskal-Wallis test, H = 4.27, df = 3, p = 0.233). Pairwise Wilcoxon comparisons also showed no significant differences between groups. Therefore, curated marker richness alone did not discriminate probiotic, potentially probiotic, neutral, and pathogenic genomes. This result indicates that the informative signal of the curated marker matrix was related mainly to marker composition and distribution, not to the total number of detected markers.

This compositional structure was supported by the Jaccard-based PCoA of curated marker profiles (Fig. 7B). The first two coordinates explained 51.93% of the total variation, with PCoA1 accounting for 41.01% and PCoA2 for 10.92%. Probiotic and potentially probiotic genomes occupied overlapping regions of the ordination space, whereas several pathogenic and neutral genomes were positioned toward distinct areas. Accordingly, PERMANOVA confirmed a significant group effect on curated marker composition (R² = 0.310, F = 6.59, p < 0.001). In parallel, beta-dispersion analysis was not significant (Fig. 7A), indicating that the observed separation was not mainly explained by unequal within-group heterogeneity. Together, these results support the use of curated marker composition as a more informative input for SPPI than marker richness alone.

**Fig. 7.**
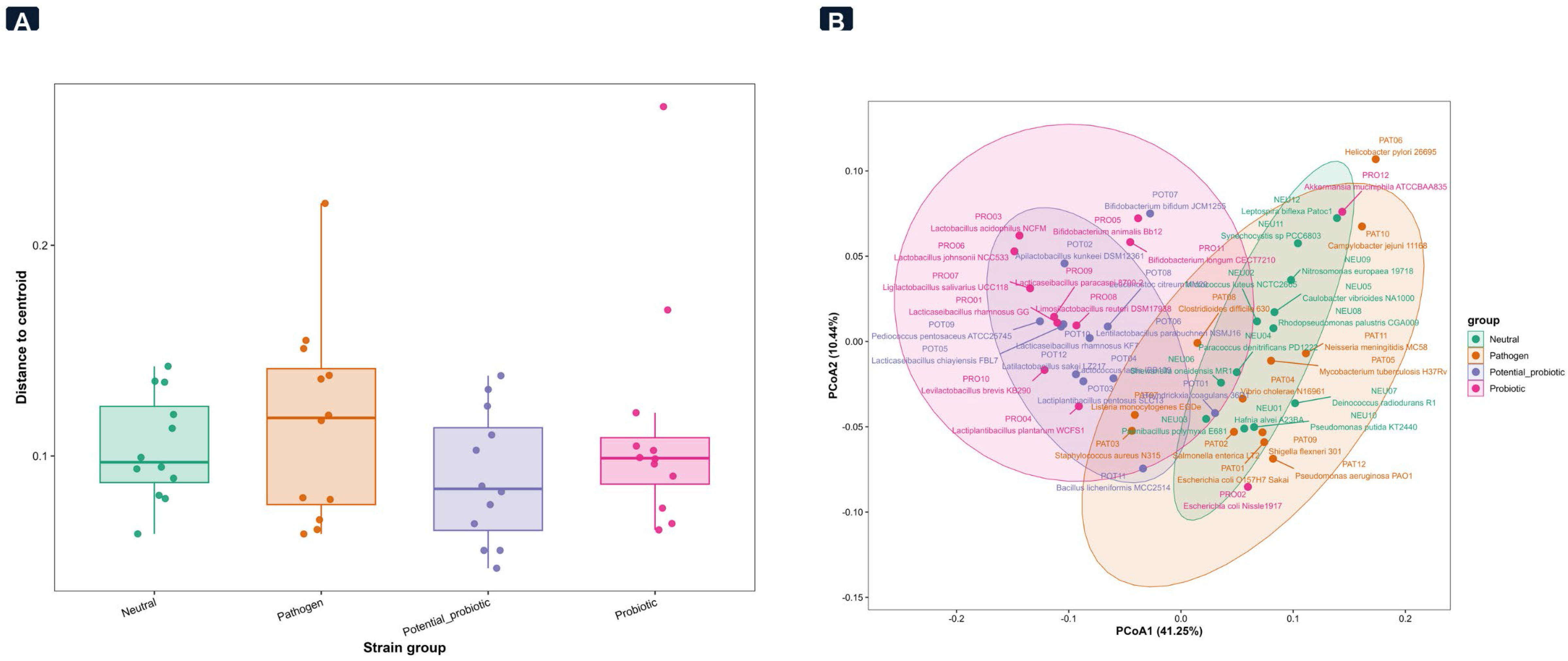
Curated probiotic marker profile structure across genome groups. A, within-group dispersion of curated marker profiles. B, multivariate structure of curated marker profiles based on Jaccard distance and principal coordinates analysis (PCoA).

### AHP-weighted SPPI 2.8 and genome prioritisation

The retained SPPI 2.8 configuration was applied after feature-level mapping and curation of the SPPI scoring inputs and represented the final upstream scoring output before FCE classification. At this stage, each genome was represented by binary scoring-feature values integrated with the final AHP-derived weights. Specifically, safety-related evidence reduced the final interpretation of a genome, whereas survival-colonisation and functional-benefit components contributed positively to the score.

Representative SPPI 2.8 outputs for the highest-scoring genome of each reference group are reported in Table 3. At the group level, the final SPPI differed significantly among the four genome groups (Kruskal-Wallis test, H = 39.69, df = 3, p < 0.001) (Fig. 8). Probiotic and potentially probiotic genomes showed close mean scores, with values of 0.895 and 0.893, respectively, and no significant difference between these two groups after correction. Neutral genomes showed an intermediate mean score of 0.829, whereas pathogenic genomes showed the lowest mean score of 0.377. Pairwise comparisons confirmed that neutral and pathogenic genomes were statistically separated from probiotic and potentially probiotic genomes, while the two favourable groups remained close.

**Fig. 8.**
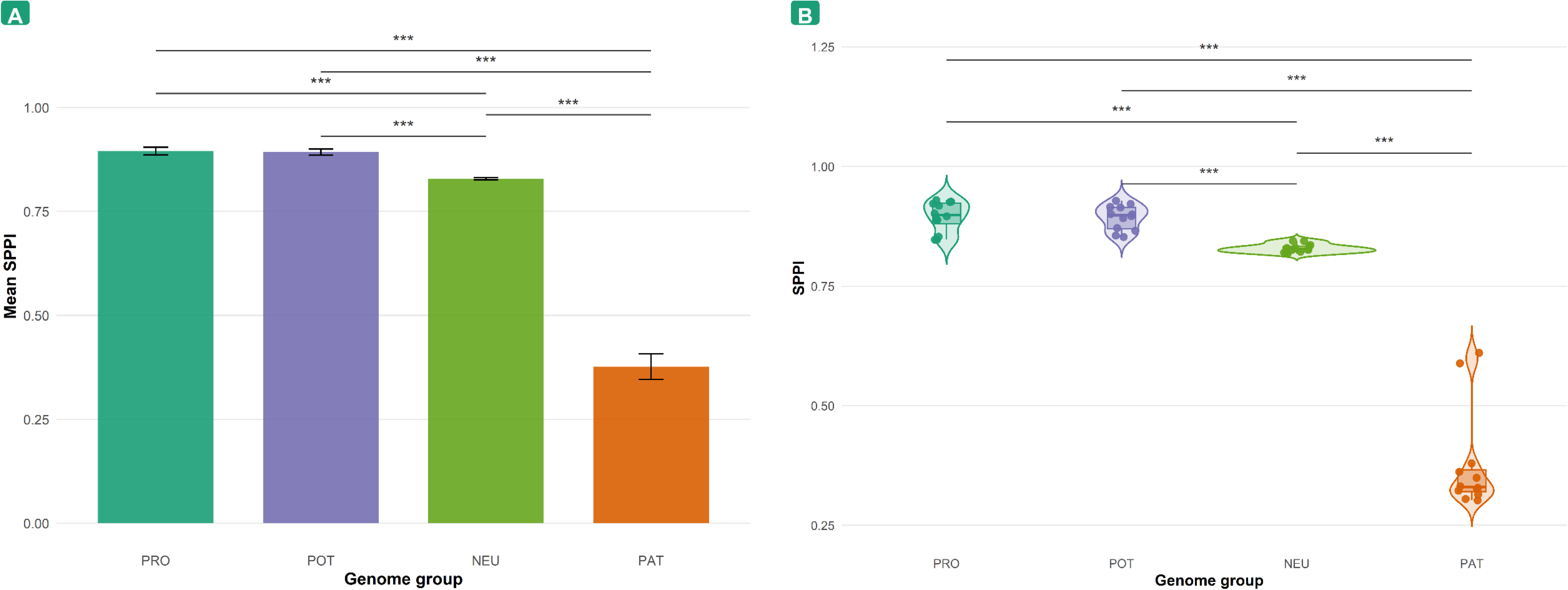
Final SPPI 2.8 distribution across the four genome groups. A, group mean of the final SPPI 2.8 values. B, individual SPPI 2.8 score distribution across the four genome groups. *: p < 0.05, **: p < 0.01, ***: p < 0.001.

**Table 3.** Representative SPPI 2.8 outputs for the highest-scoring genome of each benchmark group.

| Benchmark group | Representative genome | Genome ID | Safety | Survival-colonisation | Functional benefits | SPPI |
| --- | --- | --- | --- | --- | --- | --- |
| Probiotic | *Lactobacillus acidophilus* NCFM | PRO03 | 0.992 | 0.732 | 0.786 | 0.929 |
| Potential probiotic | *Lacticaseibacillus chiayiensis* FBL7 | POT05 | 0.990 | 0.716 | 0.808 | 0.928 |
| Neutral | *Hafnia alvei* A23BA | NEU01 | 0.991 | 0.683 | 0.112 | 0.845 |
| Pathogen | *Campylobacter jejuni* 11168 | PAT10 | 0.705 | 0.571 | 0.051 | 0.611 |

Note. Safety, Survival-colonisation, and Functional benefits correspond to pillar values obtained after subpillar-level integration of curated in silico markers using local AHP weights. The final SPPI was then obtained by multiplying these pillar values by their global AHP pillar weights

The ordered genome ranking further confirmed this separation (Fig. 9). The first 24 ranks were occupied by probiotic or potentially probiotic genomes, whereas neutral genomes started after this favourable block and pathogenic genomes occupied the lower part of the ranking. This pattern indicates that SPPI 2.8 provided a structured prioritisation of genomes within the analysed dataset. Importantly, the separation was not driven by simple marker accumulation, but by the weighted integration of safety-related constraints and favourable probiogenomic evidence according to the AHP hierarchy.

**Fig. 9.**
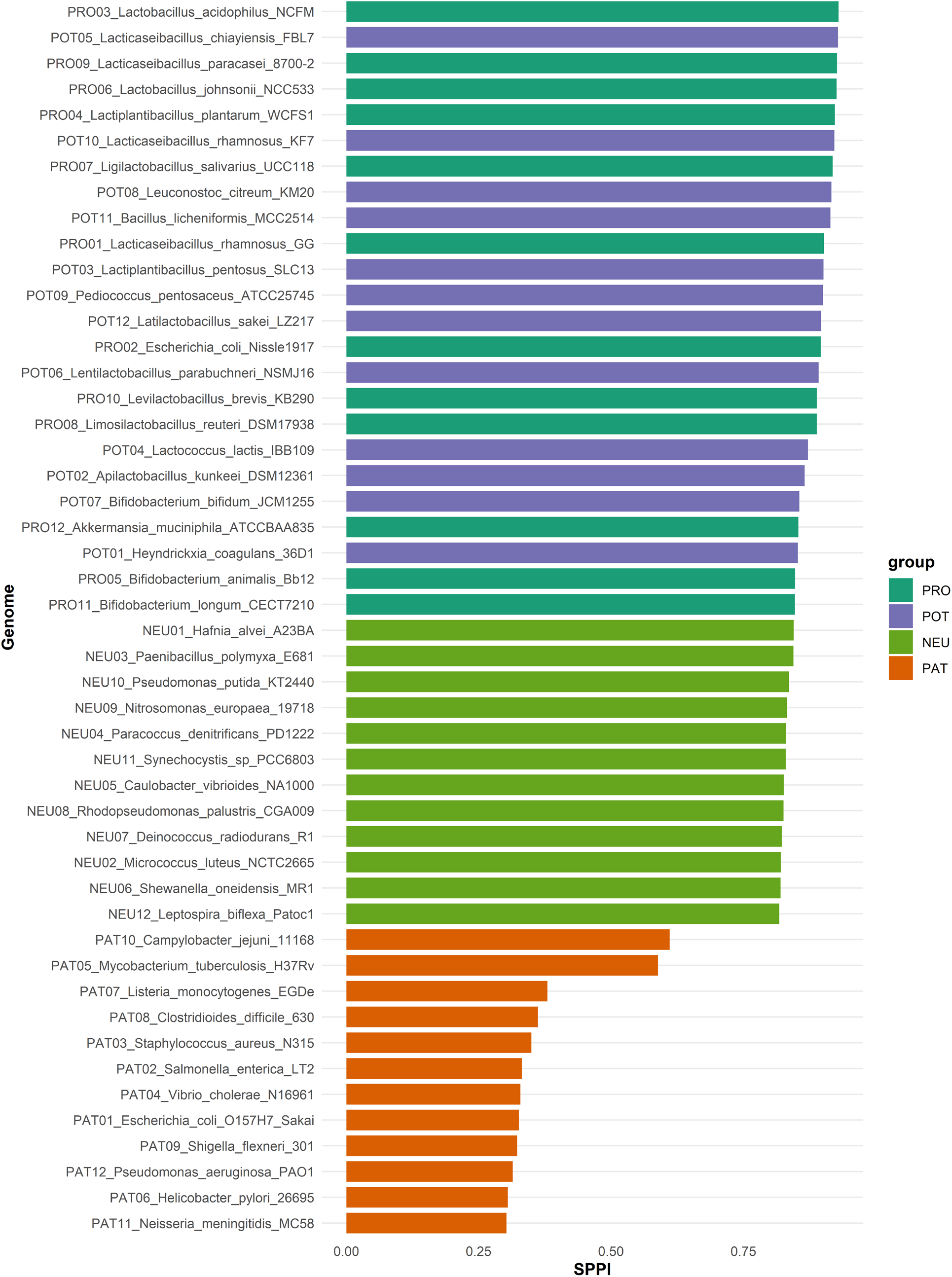
Genome ranking according to final SPPI 2.8 values. Bar plot showing the ordered SPPI 2.8 values across the 48 analysed genomes.

### Subpillar-level structure of SPPI 2.8

A standardised group mean heatmap was used to examine how the retained SPPI 2.8 subpillar scores varied across genome groups and how subpillars clustered according to their contribution profiles (Fig. 10A). The clearest risk-associated signal was observed for A2 and A3, corresponding to virulence-associated determinants and toxin-related signals. These two subpillars co-clustered and showed elevated standardised values in pathogenic genomes, whereas probiotic, potentially probiotic, and neutral genomes showed low values. In contrast, A1 and A4 did not show the same level of pathogenic enrichment, indicating that the Safety pillar was not driven uniformly by all its subcomponents.

**Fig. 10.**
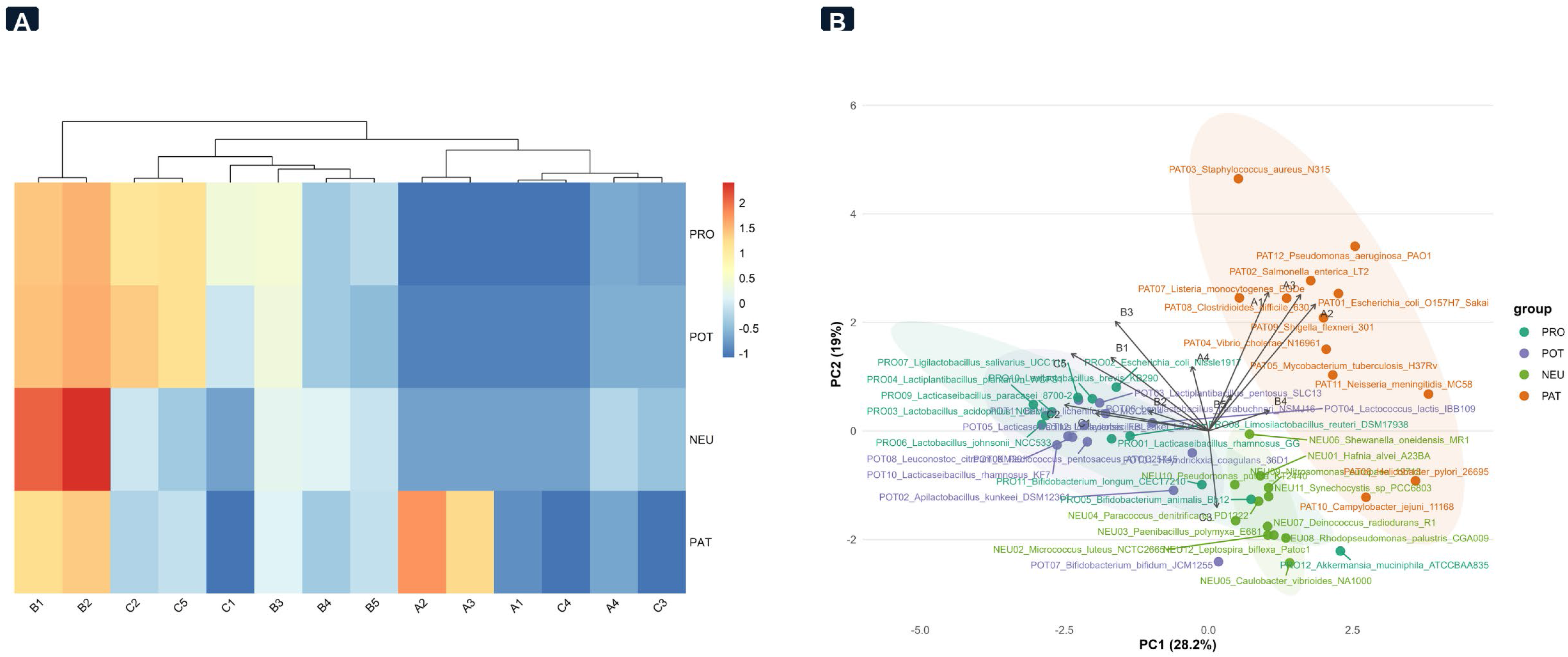
Multivariate structure of SPPI 2.8 subpillar profiles. A, standardised group mean of SPPI 2.8 subpillar scores shown as a heatmap. B, principal component analysis (PCA) based on retained subpillar scores. PC1, first principal component; PC2, second principal component; PRO, probiotic genomes; POT, potentially probiotic genomes; NEU, neutral genomes; PAT, pathogenic genomes.

Within the positive components, B1 and B2 formed a close sub-cluster and showed high values across several genome groups, including neutral genomes. This pattern indicates that acid and bile tolerance-related signals and general stress-response traits were not specific to probiotic genomes alone, but may reflect broadly distributed adaptive capacities. Other positive subpillars, particularly C2, C5, B3, and C1, contributed more directly to the differentiation of favourable profiles through beneficial metabolites, immunomodulatory potential, mucus adhesion and persistence, and antimicrobial activity.

PCA of retained subpillar scores confirmed this multivariate structure (Fig. 10B). The first two principal components explained 47.2% of the total variance, with PC1 accounting for 28.2% and PC2 for 19.0%. Probiotic and potentially probiotic genomes were mainly positioned on the negative side of PC1, whereas pathogenic genomes were located on the positive side. Neutral genomes occupied an intermediate region. PERMANOVA detected a significant group effect on subpillar score profiles (R² = 0.581, F = 20.37, p < 0.001). In the retained SPPI 2.8 configuration, C4 preserved its place in the theoretical hierarchy but was assigned a constant-zero operational value in the final scoring matrix. This choice reflected the limited group-level separation provided by the antioxidant subpillar in the analysed dataset; consequently, C4 did not contribute to PCA separation.

### FCE iteration 7.0 classification performance and sentinel genome audit

After SPPI 2.8 ranking, FCE Iteration 7.0 was applied to assign each genome to one of three final classes: Cat1, grouping probiotic and potentially probiotic genomes; Cat2, corresponding to neutral genomes; and Cat3, corresponding to pathogenic genomes. This step converted the continuous SPPI output into categorical assignments according to fuzzy membership principles (Yang et al., 2025; Zadeh, 1965).

Under leave-one-out cross-validation, all 48 genomes were assigned to their expected reference classes, including 24 Cat1 genomes, 12 Cat2 genomes, and 12 Cat3 genomes (Fig. 11A). The retained model achieved an accuracy, balanced accuracy, macro-precision, macro-recall, and macro-F1 of 1.000 within the analysed dataset. These values reflect internal classification performance on the curated 48-genome dataset and do not represent external validation on independent genomes.

**Fig. 11.**
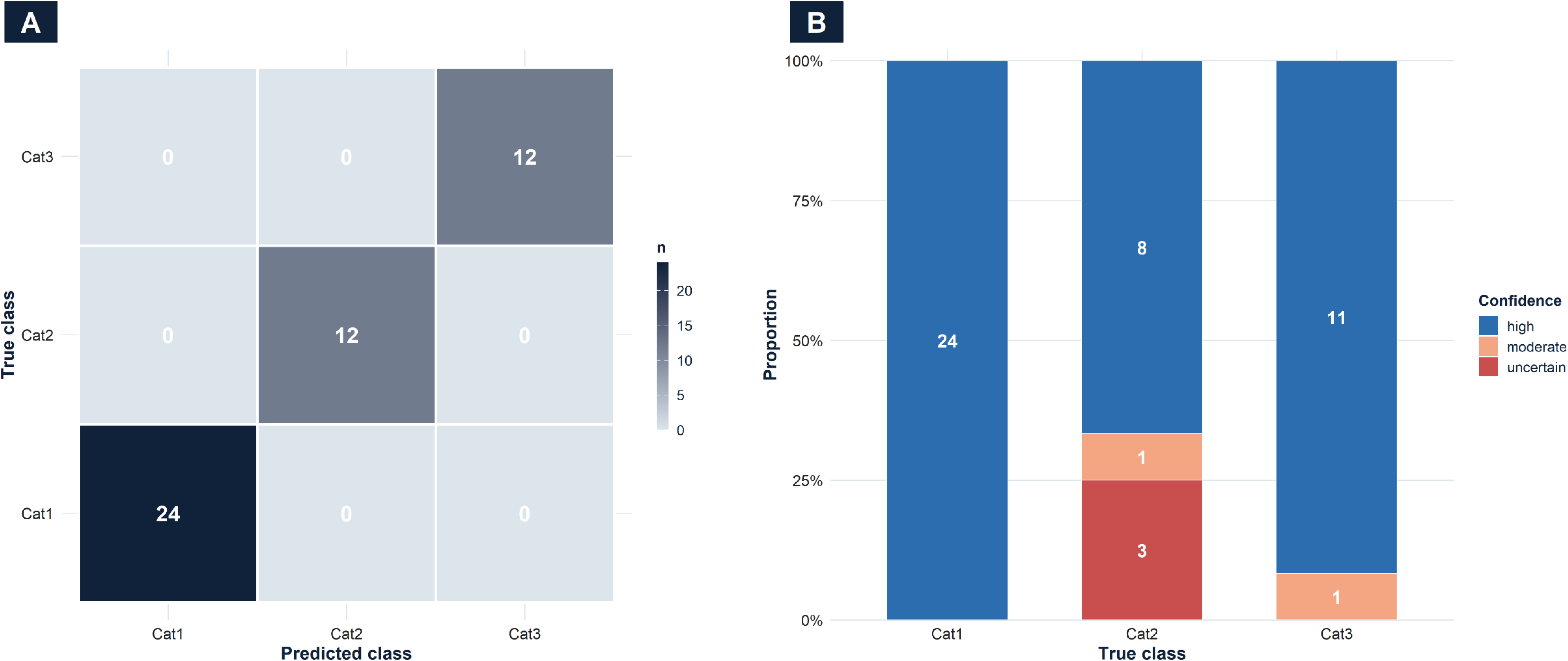
Retained FCE Iteration 7.0 classification output. A, confusion matrix of FCE Iteration 7.0 under leave-one-out cross-validation. B, confidence composition across true classes.

The confidence composition showed that Cat1 and Cat3 assignments were the most stable, whereas Cat2 remained the most borderline class (Fig. 11B). This pattern is consistent with the intermediate status of neutral genomes, which may contain broad adaptive traits without being classified as favourable probiogenomic profiles or as risk-associated pathogenic profiles.

The sentinel audit was used to examine genomes that had produced borderline or biologically complex assignments during model development (Table 4). *Escherichia coli* Nissle 1917 (PRO02) remained assigned to Cat1 despite safety-relevant genomic signals, including a colibactin-associated profile consistent with reports describing Nissle 1917 as a *pks*-positive probiotic strain (Di Pierro et al., 2026; Nougayrède et al., 2021). *Akkermansia muciniphila* ATCC BAA-835 (PRO12) was also assigned to Cat1, whereas neutral and pathogenic sentinel genomes remained assigned to their expected classes. Together, these results indicate that FCE Iteration 7.0 stabilised class assignment within the analysed dataset while preserving the separation between favourable, neutral, and risk-associated profiles.

**Table 4.** Sentinel genome classification under the retained FCE Iteration 7.0 model.

| Sentinel genome | Group | Iteration 6 classification | FCE 7.0 / SPPI 2.8 classification |
| --- | --- | --- | --- |
| PRO02_Escherichia_coli_Nissle1917 | PRO | Cat1 / uncertain | Cat1 |
| PRO12_Akkermansia_muciniphila_ATCCBAA835 | PRO | Cat2 / uncertain | Cat1 |
| NEU03_Paenibacillus_polymyxa_E681 | NEU | Cat2 / uncertain | Cat2 |
| PAT05_Mycobacterium_tuberculosis_H37Rv | PAT | Not available in retained Iteration 6 output | Cat3 |
| PAT06_Helicobacter_pylori_26695 | PAT | Cat3 / high | Cat3 |
| PAT08_Clostridioides_difficile_630 | PAT | Cat3 / high | Cat3 |

PRO, probiotic genomes; POT, potential probiotic genomes; NEU, neutral genomes; PAT, pathogenic genomes. Cat1, probiotic and potential probiotic genomes; Cat2, neutral genomes; Cat3, pathogenic genomes.

### Comparative evaluation of FCE and ProbML predictions

The retained FCE Iteration 7.0 output was compared with ProbML predictions generated on the same 48 genomes. ProbML is a machine-learning genome classifier developed to identify probiotic organisms from whole-genome sequence data (Orkkatteri Krishnan et al., 2025). For this comparison, the three FCE classes were transformed into a binary structure: Cat1 was considered probiotic, whereas Cat2 and Cat3 were considered non-probiotic.

During exploratory use of ProbML, repeated analysis of the same genome could produce different probability scores and, in some cases, different binary predictions. Therefore, the comparison reported here was based on the ProbML outputs saved at the time of analysis, including the recorded analysis date, model used, score distribution, and prediction label.

Compared with the retained FCE output, ProbML showed lower binary performance on the same 48-genome dataset (Fig. 12A). Its accuracy reached 0.854, with Cat1 recall of 0.958, non-Cat1 specificity of 0.750, Cat1 precision of 0.793, and Cat1 F1-score of 0.868. The main difference concerned non-Cat1 specificity, indicating that ProbML assigned probiotic predictions to some genomes belonging to the neutral or pathogenic groups. Discordance analysis identified seven genomes with different binary predictions between FCE and ProbML (Fig. 12B). One Cat1 genome, *Escherichia coli* Nissle 1917 (PRO02), was classified as probiotic by FCE but non-probiotic by ProbML. Conversely, six genomes were classified as non-probiotic by FCE but probiotic by ProbML, including four neutral genomes and two pathogenic genomes. Agreement analysis showed a Cohen’s kappa of 0.708, indicating substantial agreement, while McNemar’s test was not significant (p = 0.131). Thus, this comparison describes prediction behaviour on the same curated dataset, without constituting evidence of general performance on external genome collections.

**Fig. 12.**
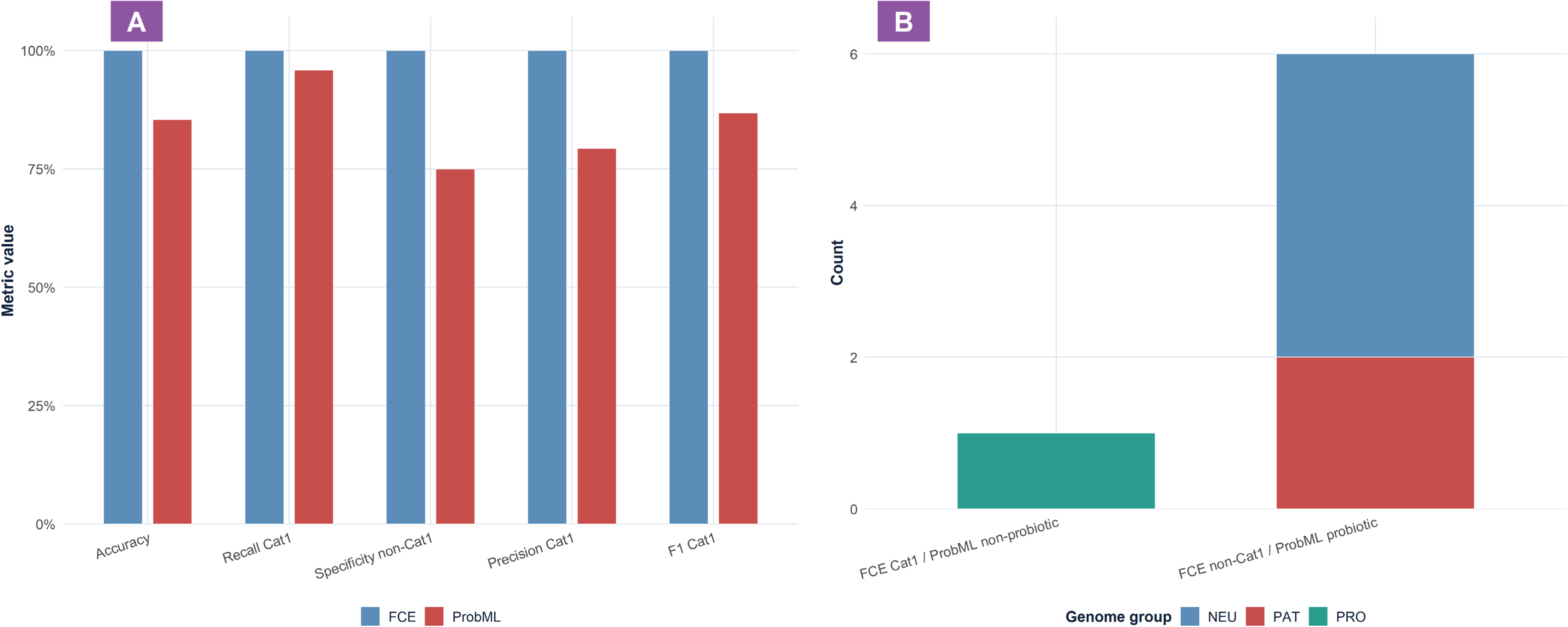
Binary comparison between FCE Iteration 7.0 and ProbML predictions. A, binary performance metrics for FCE and ProbML. B, discordant predictions according to genome group.

## Discussion

The results obtained in this study show that SPPI acts as a probiogenomic triage and prioritisation system. It does not establish that a genome belongs to a biologically confirmed probiotic strain; instead, it organises heterogeneous genome-derived evidence into a weighted interpretation that can guide early candidate selection. This distinction is central, because probiotic potential inferred from genome data remains predictive and must still be confirmed by targeted phenotypic, in vitro, and in vivo validation.

Within the analysed 48-genome dataset, SPPI 2.8 combined with FCE 7.0 produced a coherent separation between favourable, intermediate, and risk-associated genomic profiles. Probiotic and potentially probiotic genomes were positioned as favourable candidates, pathogenic genomes were shifted away from this region, and neutral genomes occupied an intermediate position. This behaviour is consistent with the intended role of the system: not to assign probiotic status directly, but to prioritise genomes according to the balance between safety constraints and supportive functional evidence.

The internal classification performance of FCE 7.0 should be interpreted within this context. The model reached complete class assignment within the analysed dataset, which indicates that the retained SPPI-FCE configuration overperformed on this curated 48-genome set. This result reflects the discriminant structure produced by the selected variables and by the two-stage classification logic. However, it should not be generalised beyond the dataset used here. The membership functions and decision rules were stabilised on the same genome collection; therefore, larger datasets, additional taxa, or genomes with more mixed profiles may require recalibration.

The behaviour of SPPI can be explained by its weighting structure. AHP assigned the dominant contribution to Safety, making the final score highly sensitive to risk-associated genomic evidence. This structure is biologically coherent, because a candidate carrying problematic safety signals should not be prioritised only because it also contains favourable functional markers. At the same time, SPPI did not function as a purely exclusion-based system. Survival-colonisation and functional-benefit components refined the ranking among genomes that did not present strong safety constraints.

This balance is important because many markers used in probiogenomic evaluation are not probiotic-specific. Survival, stress-adaptation, metabolic, and host-interaction signals can occur in probiotic, neutral, environmental, commensal, or pathogenic bacteria (Gomri et al., 2026; Neres Rodrigues et al., 2026; Papadimitriou et al., 2015). Therefore, their biological interpretation depends on their composition within the genome profile and on their relationship with safety-related evidence. In this study, total curated marker richness alone did not explain the separation between genome groups, whereas marker composition and weighted integration provided a clearer basis for prioritisation. This supports the use of a structured multicriteria approach instead of a simple presence/absence accumulation of favourable markers.

The sentinel genome audit reinforced this interpretation. Borderline or biologically complex genomes were useful for verifying that the system did not simply reward favourable signals. Genomes carrying broad adaptive traits were not automatically classified as favourable, and genomes with risk-associated profiles were not rescued by isolated positive components. This behaviour supports the idea that SPPI-FCE handled mixed genomic profiles through a balance between negative safety evidence and positive functional support.

The comparison with ProbML provided an external reference for interpreting prediction behaviour on the same dataset (Orkkatteri Krishnan et al., 2025). The objective was not to demonstrate general superiority, because ProbML and SPPI-FCE do not follow the same modelling logic. ProbML provides a binary machine-learning prediction, whereas SPPI-FCE combines weighted criteria and fuzzy class assignment. The discordant cases nevertheless showed that binary prediction and multicriteria prioritisation can lead to different interpretations, especially for neutral or pathogenic genomes carrying some favourable signals. This supports the value of a transparent system in which the contribution of safety, survival-colonisation, and functional evidence can be examined explicitly.

The expert-based weighting step also defines part of the interpretative scope of the system. Although the retained expert panel provided a relevant basis for AHP weighting, the final weights remain dependent on expert judgement and could change with a larger or differently composed panel. The consistency-based curation reduced the influence of incoherent pairwise comparisons and is consistent with controlled expert aggregation in AHP studies (Forman & Peniwati, 1998; Frish et al., 2025; Lukinskiy, Lukinskiy, & Bazhina, 2025; Salomon & Gomes, 2024). However, this does not remove the fact that AHP-derived weights represent structured expert preference rather than experimentally measured biological effects.

The dataset itself also limits the scope of interpretation. The 48-genome collection was appropriate for developing and stabilizing the scoring and classification workflow, but it cannot represent the full diversity of bacterial genomes. The choice to retain complete, high-quality, well-documented RefSeq assemblies improved traceability and screening reliability, but reduced the number of eligible strains. This is relevant because many experimentally described probiotic candidates do not yet have complete genome assemblies. ProBio-Ichnos illustrates this gap by reporting a large catalogue of probiotic strains but a much smaller number of associated genomes (Tsifintaris et al., 2024). Fragmented assemblies may also affect the detection of mobile elements, genomic islands, plasmid-associated regions, or biosynthetic loci, which can influence downstream microbial genome interpretation (Trigodet et al., 2026).

The screening outputs are also dependent on genome quality, annotation accuracy, database content, detection thresholds, and tool versions. Genomic repositories and functional databases are regularly updated, and marker interpretation may change as new annotations become available. The results therefore reflect the state of the genomes, databases, and tools used at the time of analysis. Future re-analysis will be necessary to keep SPPI aligned with updated probiogenomic resources, particularly for safety-related traits whose interpretation depends on genomic context and mobility. This point is especially relevant because mobile genetic elements remain major drivers of horizontal gene transfer and antimicrobial resistance dissemination (Hossain, 2022).

Scalability remains another methodological consideration. The local pipeline was traceable and reproducible for the curated dataset, but larger applications involving hundreds or thousands of genomes would require more efficient execution strategies, including parallelization, containerized workflows, and server or high-performance computing (HPC) resources. This is consistent with broader challenges in high-throughput microbial genomics, where computational bottlenecks, dependency management, database bias, storage requirements, and reproducible execution remain major concerns (Agudelo-Romero et al., 2025; Houmenou et al., 2025; Ismail & Amarasoma, 2025; Lacy-Roberts et al., 2026; Langer et al., 2025).

Overall, SPPI should be interpreted as a transparent decision-support layer for early probiotic candidate prioritisation. Its main contribution is to make genome-based evaluation more structured by separating safety constraints from supportive functional evidence and by converting these signals into an interpretable score and class assignment. The system can help select candidates for deeper experimental assessment, but it should not be considered a universal predictor of probiotic efficacy or a substitute for biological validation.

## Conclusion

This study developed SPPI, an AHP-guided fuzzy multicriteria system for the in silico prediction and prioritisation of probiotic potential across bacterial genomes. The approach combined literature-guided probiogenomic criteria, expert-derived AHP weights, automated genome screening outputs, weighted scoring, and downstream FCE classification into a structured decision-support workflow.

The retained SPPI 2.8 configuration organised probiotic candidate assessment around three main dimensions: Safety, Survival-colonisation, and Functional and Metabolic benefits. The weighting structure emphasised that safety should be evaluated before functional promise, while favourable survival-colonisation and functional-benefit signals should refine candidate prioritisation only in genomes with acceptable safety profiles.

Applied to the curated 48-genome dataset, SPPI enabled genome ranking according to weighted probiogenomic evidence, and FCE Iteration 7.0 converted this continuous score into operational classes corresponding to favourable, neutral, and risk-associated profiles. The comparison with ProbML further positioned the SPPI-FCE workflow against an external genome-based prediction tool on the same dataset.

Taken together, this work supports a transition from simple marker detection toward structured, weighted, and interpretable probiogenomic prioritisation. SPPI should not be considered a substitute for experimental validation, but it can serve as an early filtering layer in the probiotic selection pipeline, where risk-associated profiles are identified first, promising genomes are prioritised according to weighted evidence, and selected candidates are then directed towards targeted in vitro and in vivo validation.

Future work should apply the system to larger and more taxonomically diverse genome collections, integrate additional high-quality genomes from underexplored bioresources, and connect genomic predictions with transcriptomic, proteomic, metabolomic, and phenotypic data. In the long term, integration with artificial intelligence and machine-learning approaches could support automatic model updating, improve feature selection, and strengthen large-scale probiogenomic candidate prioritisation.

## Acknowledgements

The authors would like to thank the national and international experts who participated in the AHP survey and contributed their evaluations to this study.

## Author contributions statement

NEO: Conceptualisation, Methodology, Data curation, Formal analysis and investigation, Visualisation, Writing - original draft preparation, Writing - review and editing. MAG: Conceptualisation, Methodology, Pipeline development, SPPI/AHP calculator development, Validation, Formal analysis and investigation, Visualisation, Writing - review and editing, Supervision. MEHEO: Conceptualisation, Methodology, Validation, Writing - review and editing

## Ethics approval and consent to participate

Not applicable.

## Data availability

The datasets and computational materials supporting this study, including genome metadata, curated marker tables, AHP outputs, SPPI 2.8 calculation files, FCE 7.0 classification outputs, ProbML comparison files, statistical analysis outputs, figures, tables, and source scripts, are available in the Zenodo repository associated with this study: https://doi.org/10.5281/zenodo.21446055

